# Mechanistic Model of Replication Fork Progression in Yeast

**DOI:** 10.1101/2021.10.03.462908

**Authors:** Ghanendra Singh

## Abstract

Replication fork progression complex plays an essential role during DNA replication. It travels along with the DNA with a particular speed called replication fork speed. Faithful duplication of the genome requires strict control over replication fork speed. Both acceleration and pausing mechanisms of the replication fork complex are regulated at the molecular level. Based on the experimental evidence, DNA replicates faster in normal cells than cancer cells, whereas cancer cells duplicate themselves more quickly than normal cells. Then in principle, accelerating the replication fork complex in cancer cells beyond a specific threshold speed limit can cause DNA damage and plausibly kill them. A modular mathematical model is proposed to explain the dynamics of replication fork control during DNA replication using the underlying molecular mechanisms in yeast which can extend to the mammalian system in the future.

## Introduction

The rate of DNA replication fork movement changes during the S phase in mammalian cells. More replication origins fire during fast DNA replication. The rate of fork movement increases by about threefold from the early S phase to the later S phase, with the most dramatic change occurring in the first hour of the S phase [13]. Changes in the replication fork rate account for the variation in S phase duration of the yeast *S.cerevisiae* [26]. Replication fork speed reduces while approaching stalling or commonly known as replication pausing sites in yeast [6]. Correlation between inter origin distance and fork velocities exists [8]. Higher no. of origin firing or small origin to origin distance results in slow replication fork rate. Less no. of origin firing or large origin to origin distances results in higher replication fork rates. A hypothesis was proposed by [8] about the existence of a molecular mechanism for replication forks to adjust their speed during the S phase. In the first half of the S phase, gene-rich euchromatin replicates in which the density of replication origins increases by two folds. In the second half of the S phase, gene-poor heterochromatin gets duplicated. The replication origins density increases by ten folds [12]. Another experimental evidence [17] showed that the MTC (Mrc1-Tof1-Csm3) complex interacts dynamically with the replisome. Mrc1 transiently interacts with the Tof1-Csm3 complex to speed up the replication fork. Speed increased up to 2 to 3 folds which match with the earlier shown results [13].

Therefore, there might be a plausible mechanism for the replication fork speed to the timing of the S phase of DNA replication. During fork progression, when a fork approaches another incoming fork, distance between them reduces and fork speed increases which helps in completion of DNA replication in timely manner [35]. Mec1 kinase is recruited to sites of replication forks. It phosphorylates a component of the DNA replication complex, Mrc1, thereby setting up a solid-state Rad53 activation platform to initiate the checkpoint response [25]. Depletion of nucleotide signals replication stress through Mrc1 pathway which dynamically interacts with replication progression complex that results in pausing of fork [32]. Rad53 regulates replication fork restart after DNA damage in *S.cerevisiae.* A similar kind of system in E Coli was named Deadman switch [19].

However, Tof1 with Csm3 (checkpoint mediator proteins) proposed to act in pausing the Rrm3 helicase, which itself can overcome fork pausing. [22]. The protection against trapped topoisomerases conferred by the Tof1-Csm3 complex appears to require additional input from Mrc1 to promote the DNA replication checkpoint. Cdc13 binds telomerase through interaction with Est1 [9]. The role of the fork protection complex in fork acceleration and pausing is evident. But it is still unclear exactly how the two impart effects on the replisome and how they are interconnected. Recently two models postulated by [29]. The first is “pausing-centric,” and the second is “acceleration-centric” to explain fork rate control by fork protection complex.

Replication fork acceleration and pausing is an important mechanism that results in slower or faster DNA replication. Researchers experimentally showed about the acceleration or slowed down of replication fork. It’s a new approach to the cancer problem in which DNA replication is slower than normal cells, whereas cancer cells duplicate faster than normal cells [30]. There exists a need to model the mechanism of replication fork speed control operating at the molecular level. Then in principle, if we can accelerate or apply brakes to the progression of replication fork during DNA replication using external agents beyond a certain speed threshold which fork progression complex cannot handle may lead to the crash of cancer cells. The importance of investigating dynamic interaction between fork progression complex and replisome in vivo and how this process is regulated and its functional role recently featured [29].

We look at the replication fork progression mechanism as a combination of modular independent circuits connected together giving rise to this complex behavior in *S. cerevisiae.* Here, we developed the following simple mathematical models based on the underlying molecular mechanism: (1) Replication fork accelerator model-the control mechanism of replication fork speed based on S phase DNA replication checkpoint. (2) Replication fork with cell cycle CDK and DDK kinases. (3) Replication fork braking molecular mechanisms. (4) Molecular accelerator and braking mechanisms together. (5) Replication fork regulation using dNTPs (deoxyribonucleotide triphosphate). (6) Replication fork near telomeres. Later, we propose a block diagram of the complete replication fork model.

## Methods

Python is used for numerical simulation of systems of ODEs related to mathematical models. Experimental data for slow replication fork speed is taken from [30]. Another experimental data of Mrc1 dynamic interaction with Tof1 and Csm3 belonging to MTC (Mrc1, Tof1, Csm3) complex resulting in change of replication fork speed is taken from [17].

## Results

Initially, all the replication fork model components combined in a modular fashion are present in figure 1 to get an overall picture. In later sections, we visit each module independently, indicate its behavior, and show the nature of the dynamics of modules joined together.

**Figure 1:**
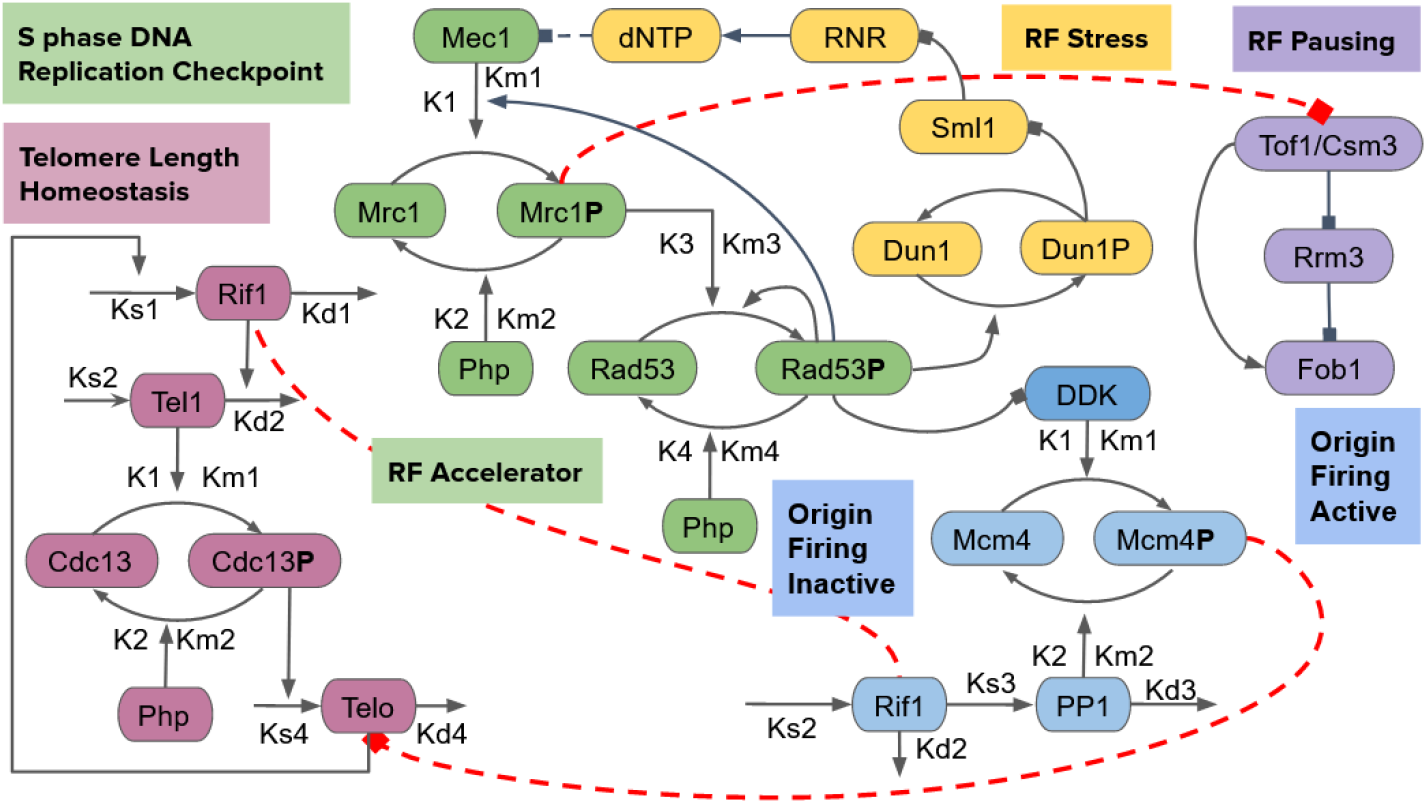
Replication Fork Model (RFM) modular circuit diagram. Different independent modules like replication fork accelerator model (in green), replication fork braking model (in violet), fork progression regulation using dNTP (in yellow), telomere length homeostasis model (in purple), and regulation by cell cycle DDK kinase (blue) are connected together to explain global fork dynamics. Red dashed lines represents inhibition mechanism except Rif1 joined with red dashed line in telomeric (in purple) and non telomeric regions (in blue).

**Figure 2:**
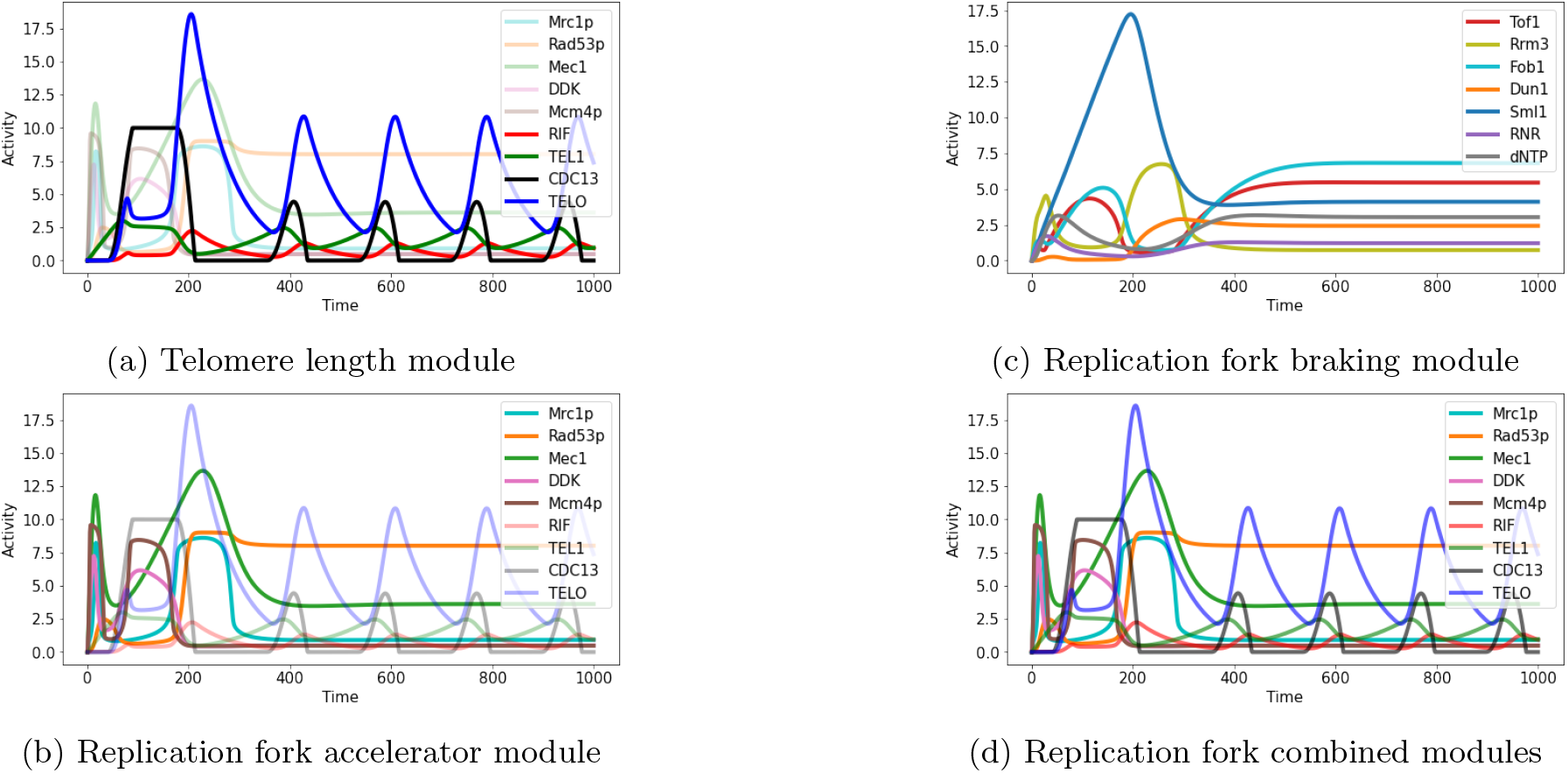
Replication Fork Model (RFM) simulation results of individual modular components. (a) Oscillatory dynamics of Rif1 (in red) and telomerase TELO (in blue) in telomere length homeostasis model. (b) Mrc1P (in light blue) and Rad53P (in orange) activation during high replication stress signal sensed by Mec1 (in green) inhibiting high DDK (in pink) activity and suppressing origin firing by dephosphorylation of Mcm4P (in brown) by PP1 during S phase replication checkpoint in fork accelerator model (c) High Mrc1P inhibits Tof1 (in red) activating Rrm3 (in yellow) leading to down regulation of Fob1 (in light blue) directly by Tof1 and indirectly by Rrm3 representing a coherent feed forward loop in fork braking model. Other factor Dun1 (in orange), Sml1 (in dark blue), RNR (in purple), and dNTP (in black) represent fork regulation by dNTP. (d) Dynamics of all modules connected together. All the parameter values are available in the supplementary material.

### Replication Fork Accelerator Model

During replication stress, Mec1 kinase protein acting as a sensor is recruited, which phosphorylates a DNA replication complex component Mrc1. Mrc1 transduces this replication stress signal to activate Rad53 [1]. Rad53 positively autoregulates itself and further mediates the phosphorylation of Mrc1. This Mrc1 mediated Rad53 hyper-phosphorylation gives rise to an immediate sensitive action of sensing any replication stress and bringing the replication fork’s passage to a halt. Rad53 in *S.cerevisiae* plays a vital role in the cellular response to DNA damage and replication checkpoint. It also activated downstream kinase Dun1p. Rad53 later inhibits cell cycle checkpoint kinase DDK and Sld3, which inhibit replication origins from firing. By the time Rad53 gets activated, Mrc1 activation sends a signal to stop the replication fork. Mrc1 directly stimulates replisome rate and is aided by Csm3/Tof1 complex.

A positive feedback loop exists in this replication stress signal transduction network of *Mec*1 → *Mrc*1 → *Rad*53 → *Mrc*1. It is unknown whether fork acceleration or pausing is regulated during the S phase, under replication stress, or in cells experiencing DNA damage. To test the hypothesis, we designed a circuit diagram given in the figure 3a to capture the dynamics of replication fork movement during the S phase. Next, we construct a simple model to explain the role of positive feedback in the DNA replication checkpoint pathway. The model is given below by the following system of equations.

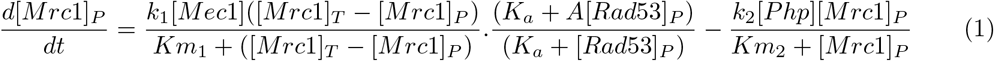

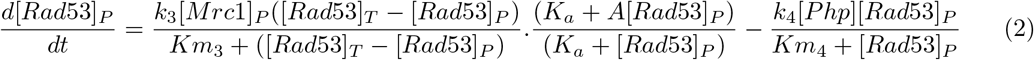

**Figure 3:**
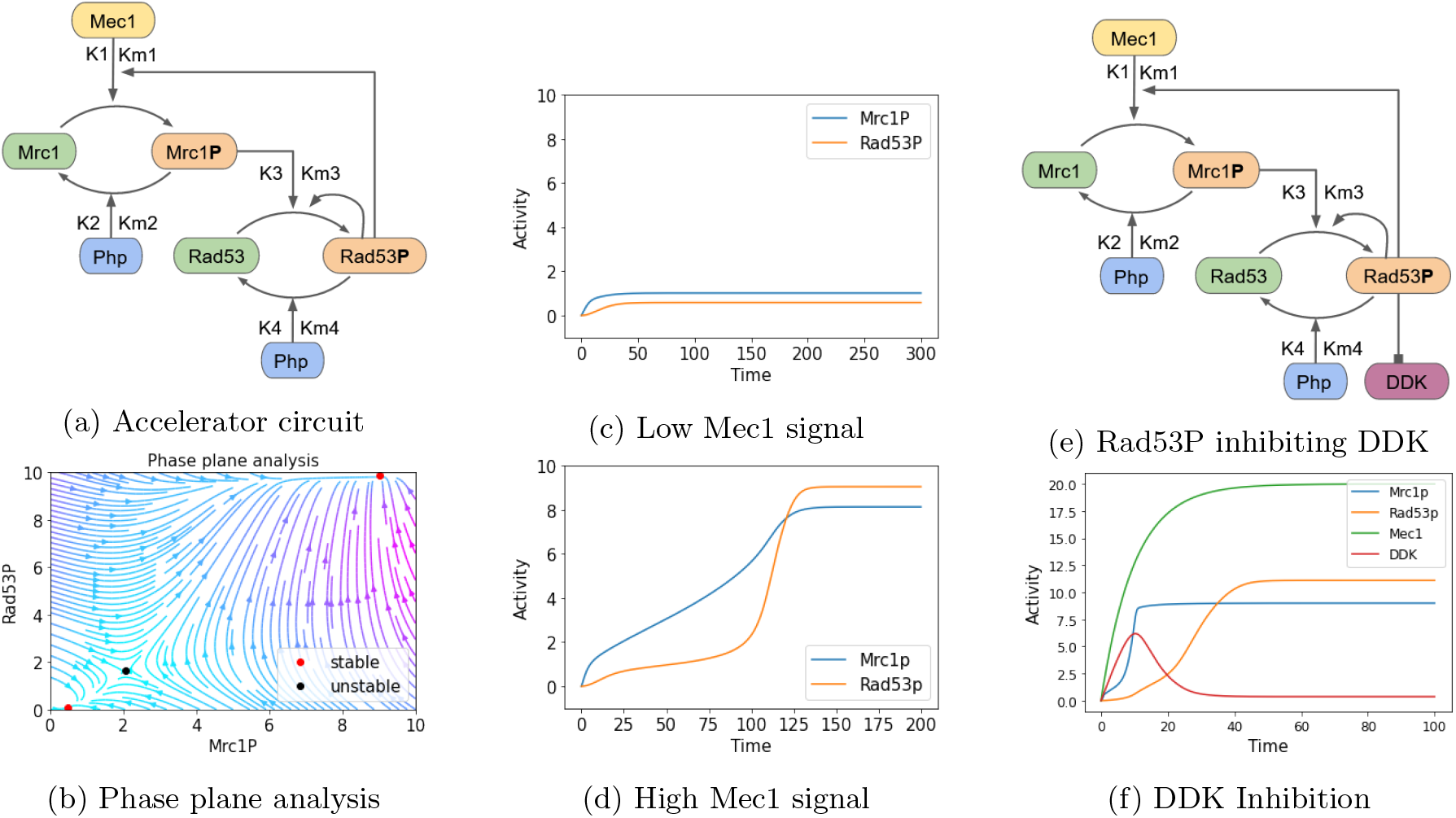
Replication Fork Accelerator Model (*a*) Biological circuit of S phase DNA replication checkpoint signaling cascade which consists of a positive feedback mechanism. Parameters for the model are: *k*_1_ = 0.05, *k*_2_ = 0.03, *k*_3_ = 0.03, *k*_4_ = 0.02, *Km*_1_ = *Km*_2_ = *Km*_3_ = *Km*_4_ = 1.0, *k_a_* = 2, *k_d_* = 0.1, *k*_*a*2_ = 0.8, and *k*_*d*2_ = 0.2. (*b*) Phase plane analysis reveals existence of two stable fixed points shown by red and an unstable fixed point shown by black. Positive feedback gives rise to bi-stability. (*c*) Mrc1 is inactive at low input Mec1(t=0) = 2. (*d*) Mrc1 gets active at high input Mec1(t=0) = 6. (*e*) Addition of DDK inhibition by Rad53 to the biological circuit diagram given in (a). (*f*) Response of the simulation results which indicates inhibition of DDK activity by sufficient Rad53P.

In equation 1, *k*_1_ is [Mec1] kinase mediated [Mrc1] phosphorylation rate and *k*_2_ is [Php] phosphatase mediated [Mrc1] dephosphorylation rate. Further in equation 2, *k*_3_ is [Mrc1] mediated [Rad53] phosphorylation rate and *k*_4_ is [Php] phosphatase mediated [Rad53] dephosphorylation rate. *Km*_1_, *Km*_2_, *Km*_3_ and *Km*_4_ are Michaelis rate constants in equation 1 and 2 respectively. [*Mrc*1_*T*_] and [*Rad*53_*T*_] total concentrations are conserved. [*Mrc*1] + [*Mrc*1_*P*_] = [*Mrc*1_*T*_] and [*Rad*53] + [*Rad*53_*P*_] = [*Rad53*_P_].

In figure 3b, phase plane analysis simulation predicts two steady, stable states. It indicates that a replication fork moves with speed regulated within a particular range near a fixed point where Mrc1 and Rad53 levels are low under normal conditions, promoting fork stability as shown in figure 3c. In this state, the replication fork speed can be controlled and varied from 1.5x to 3x times the average rate as shown by the experimental results about dynamic interaction of Mrc1 with replisome [17]. But when the replication stress arises beyond a certain threshold, as shown in figure 3d, the system transitions to another state, reducing the fork speed or pausing or halting the replication fork complex to maintain genome integrity and deal with stress factors.

### Replication Fork Rate

Replication fork progression occurs at a certain speed which varies during DNA replication. Here, we provide an expression initially for fork acceleration *f_a_* with units (*kb/min*^2^) which is rate of change of fork velocity v (*kb/min*), given by

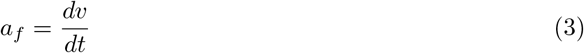

Expression for fork rate v (kb/min) is given by

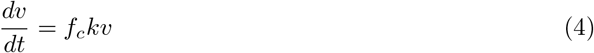

where *f_c_* is fork coefficient taken as −1 when fork rate switches from a higher to a lower value and k is a dimensionless parameter represent changes in molecular concentration of factors regulating replication fork or other barriers that modify the fork rates. Here k represents the product of ratios of Mrc1 and Rad53, as both contribute in changing the fork progression rate. Therefore, k is given by

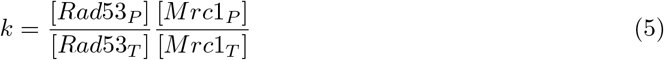

Solution of the equation 4 is given by

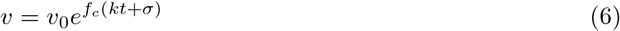

Here, initial fork velocity *v*_0_ at time (t=0) is chosen at random between (0.6 - 3.6) (kb/min) depending on the experimental data. Fork rate varies during DNA replication, hence a parameter sigma *σ* ϵ N(0,v) taken from normal distribution is chosen to represent the stochasticity in fork rate.

Putting the solution 6 in equation 3, fork acceleration is given by

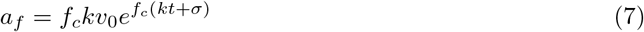

and final expression of fork rate v(kb/min) by replacing k with its value is given below.

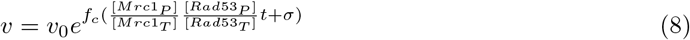

Replication of telomeres takes place at the end of the S phase when a replication origin fires near telomeric or subtelomeric regions or from a distant origins and a replication fork travels towards the chromosome end and replicates the telomeres. Assuming a single origin fires near subtelomeric DNA and a single fork progresses towards the telomere, we consider fork velocity v (kb/min) as a rate of change of DNA replication. For telomeric DNA replication, its fair to consider v as rate of change of telomere length (L). Therefore fork velocity v is given by

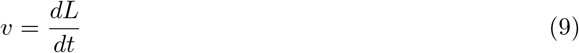

where L is telomere length and the solution of above equation 9 is given by

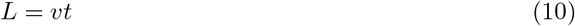

where (L) is the length of telomeric DNA replicated equal to product of fork velocity (v) and time (t). Telomere length (L) can be rewritten as total telomere length *L_t_* as sum of unreplicated telomere length *L_u_* and replicated telomere length *L_r_*. *L_t_* = *L_u_* + *L_r_*. Length of replicated and unreplicated telomere can be determined at each time step of fork progression during DNA replication.

### Role of DDK in Replication fork progression

Dbf4-dependent kinase (DDK) plays a role in replication origin firing. DDK was hypothesized to regulate the Tof1-CMG (Cdc45-MCM4-7-GINS) interaction [5]. Does the fork progression complex utilize DDK kinase to modulate fork speed? Recently identified PRDX2 with mammalian TIMELESS protein forms a replisome-based redox sensor which is involved in fork speed modulation [30]. It provides another precedent for adjusting fork speed. Additional system of equations for DDK and Mec1 is given by

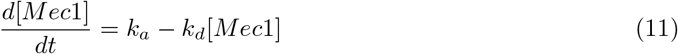

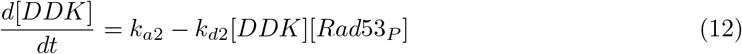

In equation 11, *k_a_* is Mec1 production rate and *k_d_* is Mec1 degradation rate. Further in equation 12, *k*_*a*2_ is DDK production rate and *k*_*d*2_ is Rad53P mediated DDK degradation rate. Cell cycle checkpoint kinases (CDK and DDK) regulates DNA replication origins. To model this effect DDK is added initially as shown in the figure 4e which is negatively regulated by Rad53 during replication stress shown by equation 12. Earlier we have considered Mec1 concentration as constant and now we consider it as a function of time (t) shown by equation 11. By adding the role of DDK and Mec1 the system of ODE is given by equations 1, 2, 11 and 12. Simulation results of these system of ODEs in figure 4f shows that starting at time (t=0) after some time when the Mrc1 mediated Mec1 dependent Rad53 concentration reaches a sufficient level it inhibits DDK resulting in down regulation of DDK activity. These system of equations shows key role of Rad53 and DDK in regulating firing of replication origins when replication stress occurs during DNA replication.

**Figure 4:**
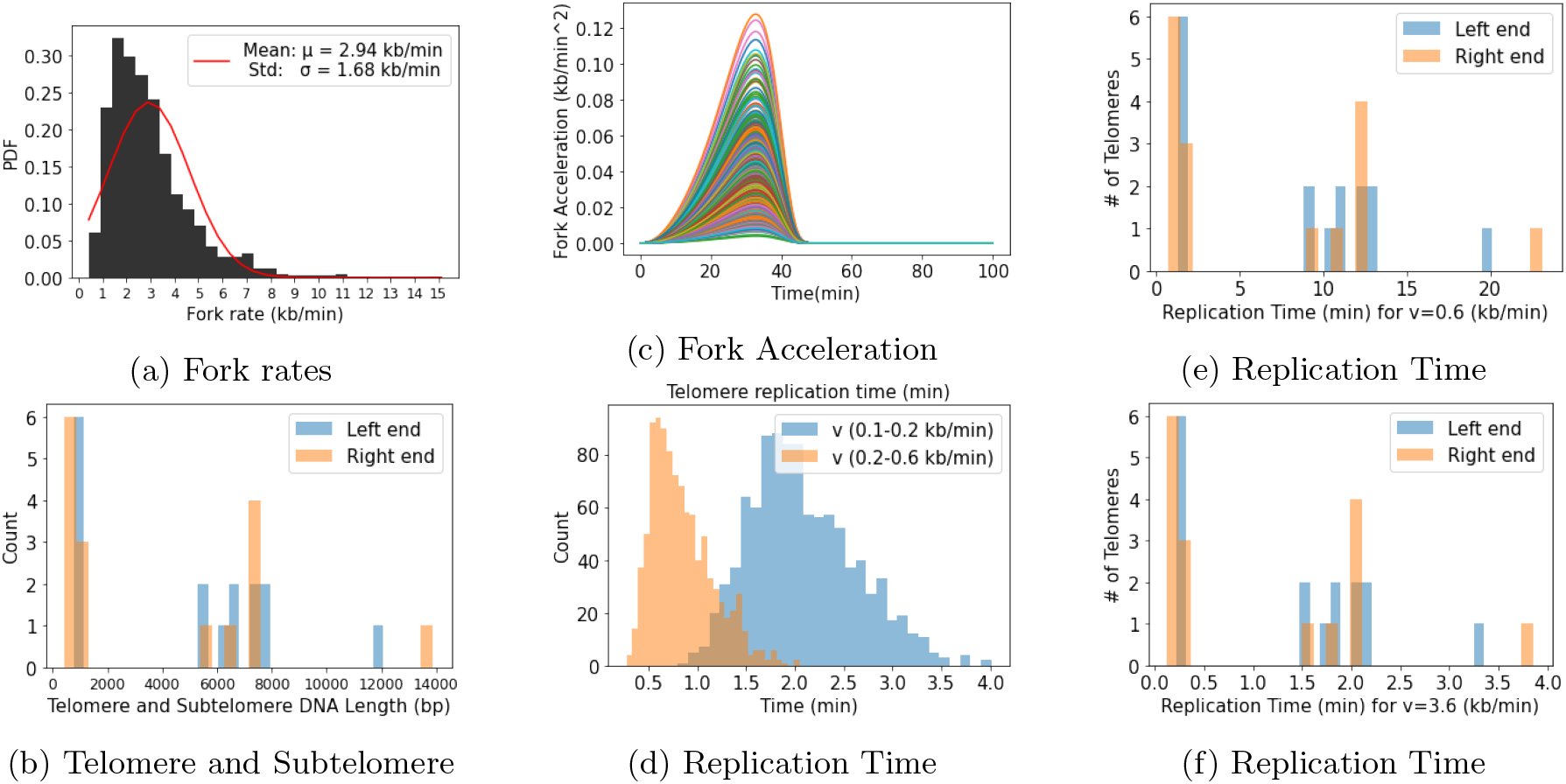
Replication Fork Speed/Acceleration dynamics (a) Distribution of fork rate v(kb/min) obtained after 1000 iterations using equation 6. (b) Left and right Telomereic subtelomeric length histogram plots based on SGD database [7]. (c) Fork acceleration *f_a_* for 1000 iteration of transition in fork velocity from higher to lower value during DNA replication stress. (d) Mimicing telomere length distribution from Teixeira data to demonstrate replication time at different slow fork velocities, (e) Telomere replication time at slow fork velocity v=0.6 (kb/min). (f) Telomere replication time at fast fork velocity v=3.6 (kb/min).

### Replication Fork with Cell Cycle

We have used a simplified 2D Model of cell cycle from [10] to model the dynamics of CDK and APC to investigate its role in recruiting DNA replication complexes to fire replication origins and start fork progression. System of equations is given below.

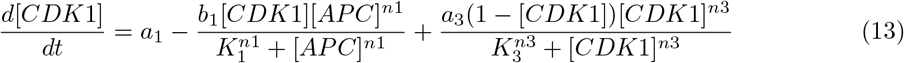

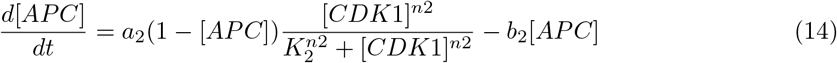

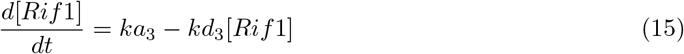

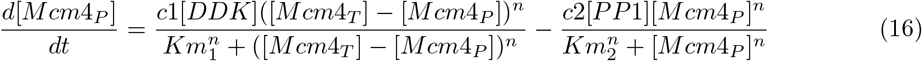

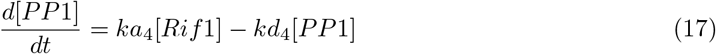

During the S phase of the cell cycle, the formation of the replication fork takes place at replication origins, and origins fire for the fork traversal. DDK phosphorylate Mcm4 subunit of the MCM2-7 complex to activate origin firing as shown in figure 5a. On the other hand, to suppress origin firing, PP1 is recruited by Rif1, which reverses the DDK-mediated phosphorylation of Mcm4. Dephosphorylation of Mcm4 results in deactivation of origin firing [20].

**Figure 5:**
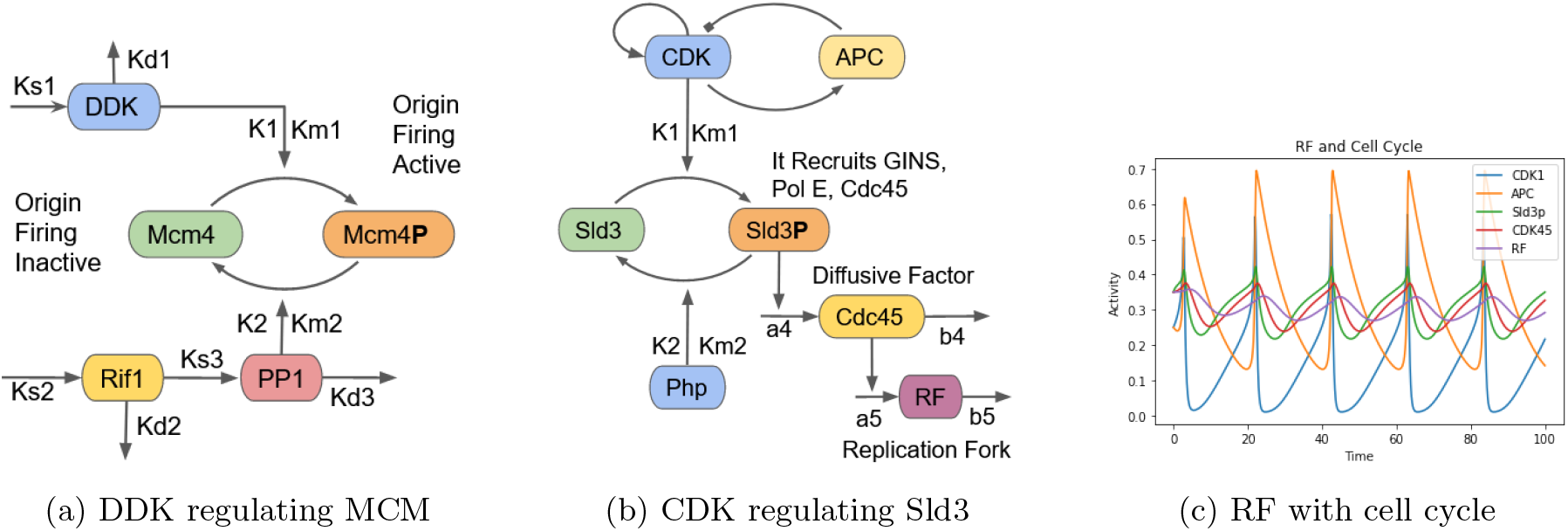
Replication Fork Model with CDK and DDK regulation (a) Initially origins which are not fired yet are considered as inactive origin. DDK kinase phosphorylates Mcm4 component of replication fork complex and on phosphorylation, origin firing becomes active and replication fork can initiate. During replication stress, Rif1 dephosphorylates Mcm4P mediated by PP1 phosphatase. (b) CDK kinase phosphorylates Sld3 which hels in recruiting GINS, Polymerase ϵ, Cdc45 diffusive factor which associate with replication origins during M phase and DDK activates replication fork during S phase. (c) Simulation of replication fork along with cell cycle kinases CDK1 (in blue) and APC (in orange). Replication fork RF (in purple) density oscillates at each cell division indicating initially low at start of S phase, maximum at mid S phase and gets low again at end of S phase.

**Figure 6:**
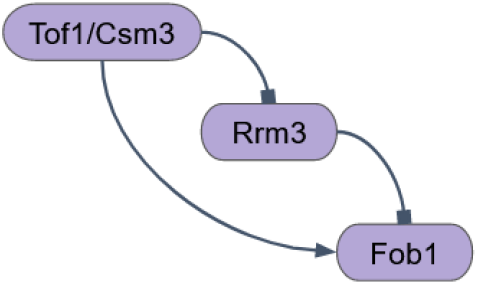
Replication Fork molecular braking mechanism. Coherent feedforward regulation of Fob1 by Tof1/Csm3. Parameters for the model are *kp*_1_ = 0.1, *bd*_1_ = 0.02, *kp*_2_ = 0.4, *bd*_2_ = 0.1, *kp*_3_ = 0.2 and *bd*_3_ = 0.04 with initial conditions Tof1(t=0) = 0, Rrm3(t=0) = 0, Fob1(t=0) = 0. Tof1/Csm3 complex positively regulates Fob1 and negatively regulates Fob1 mediated via Rrm3. Tof1 activates Fob1 directly indicating positive feed forward regulation and inhibits Rrm3 which in turn inhibits Fob1 in indirect manner indicating double negative feed forward regulation.

Equations 13 and 14 are 2D cell cycle equations from [10]. DNA replication is regulated by the cell cycle kinases, particularly during origin firing. At the end of G1 phase, CDK phosphorylate Sld2 and Sld3 as shown in 5b. Phosphorylation of Sld2 leads to the recruitment of Dbp11, GINS, and Pol(E) [24] forming a subcomplex to the replication origin throughout the cell cycle. Sld2, GINS, and Pol(E) are omitted in the figure for simplicity. DDK is active only during the S phase because Dbf4 is only expressed in late G1, and Dbf4 is degraded by APC [33]. Additional phosphorylation of MCM2-7 complex by DDK results in the arrival of GINS at the pre recognition complex results in the formation of Cdc45-MCM4-7-GINS (CMG) complex, which represents the active helicase [3] and results in the unwinding of DNA and initiates DNA replication.

In figure 5c simulation results shows cell-cycle oscillations of CDK1 and APC. Similarly, the activity of Sld3 is high at the transition of the G1/S phase of the cell cycle, indicating origins get activated. CDK45 is an important replication factor that triggers fork activation and bidirectional movement. RF is represented together as a whole replication fork complex, and its activity is initially low at the start of the S phase, which becomes high during the mid-S phase and drops down at the end of the S phase.

### Replication Fork Braking Mechanism

To examine the dynamics of DNA replication and measure the firing of replication origin timing, we normally use external agent hydroxyurea (HU). It inhibits ribonucleotide reductase (Rnr), resulting in dNTP (deoxyribonucleotide triphosphate) depletion and causing a slowdown of replication fork resulting in slow DNA replication [2]. Loss of Mrc1 gene function in yeast [16] or of Timeless in mammalian cells [30] causes slow fork progression. Replication fork pausing requires a specific fork blocking protein Fob1 to get recruited to replication fork barrier sites. Many other factors like DNA secondary structures, DNA bound proteins, etc., impede fork progression. Rrm3 is a negative regulator of fork pausing that promotes fork progression through various impediments. A conserved complex consisting of Tof1 and Csm3 in budding yeast and Swi1 and Swi3 in fission yeast plays a primary role in fork pausing [22]. Fork protection complex inhibits the activity of Rrm3 activity that removes barriers [22], and they function independently of each other [28]. We anticipate the possibility of a feedforward loop between these factors. Here, we show that the stability of the RF braking mechanism comes from coherent feedforward regulation of Fob1 by Tof1/Csm3, which is isolated from the more complex regulatory machinery of the replication fork. Tight regulation of Tof1 phosphorylation allows rapid removal of barriers. We expect coherent feedforward regulation implemented at the braking mechanism. Two distinct mechanisms activate fob1: Tof1 directly regulates Fob1 and indirectly through Rrm3. As described above, coherent feedforward regulation of Fob1 by Tof1/Csm3 plausibly intertwines with cell cycle S phase DNA replication checkpoint. Can this analysis provide insight into the regulation of RF rates? We expect this network is critical for controlling reversible fork transitions.

To understand how the Tof1/Csm3 induced Rrm3 pathway fulfills the demand of fork pausing and removing barriers, we decided to model Rrm3 feedforward regulation using differential equations. The following equations give the model:

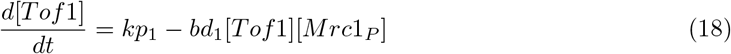

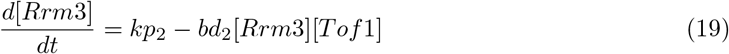

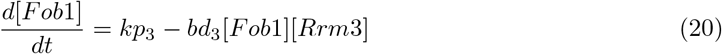

In equation 18, *kp*_1_ is [Tof1] production rate and *bd*_1_ is [*Mrc*1_*P*_] mediated [*Tof*1] degradation rate. Next, in equation 19, *kp*_2_ is [Rrm3] production rate and *bd*_2_ is [Tofi] mediated [Rrm3] degradation rate. Further in equation 20, *kp*_3_ is [Fob1] production rate and *kp*_3_ is [Rrm3] mediated [Fob1] degradation rate.

### Replication Fork Accelerator and Braking Model

Fork Speed Regulatory Network (FSRN) which regulates the speed of replication forks was proposed by [21]. Another sTOP “slowing down with topoisomerases” model was proposed [28] in which a conserved pair of budding yeast replisome components Tof1–Csm3 (fission yeast Swi1–Swi3 and human Timeless–Tipin) act as a “molecular brake” and promote fork slowdown at replication fork barriers sites, while Rrm3 assists replisome in removing protein barriers. Here, combined dynamic circuit of replication fork accelerator and braking mechanism is given in figure 7 and the simulation results shown in figure 8.

**Figure 7:**
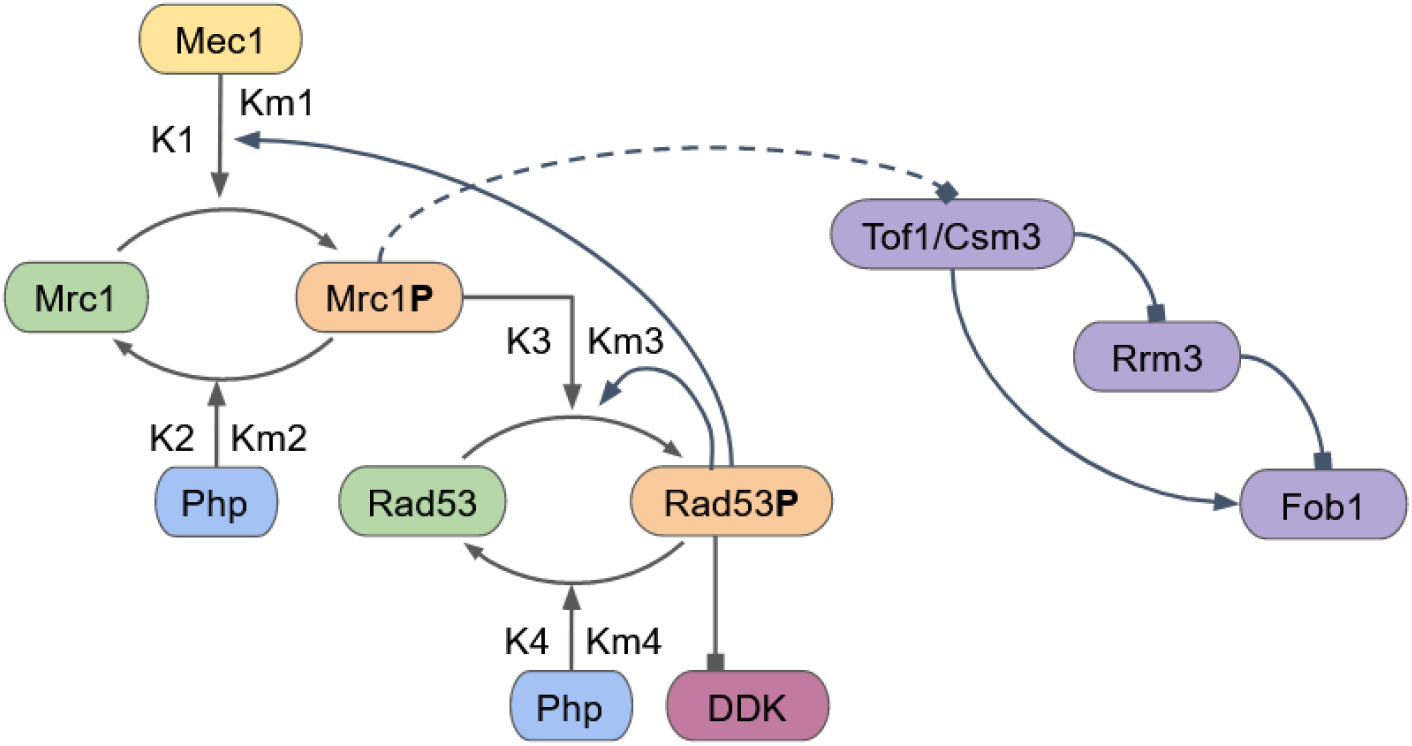
Replication Fork molecular accelerator and braking mechanism. Mrc1P interacts dynamically with Tof1/Csm3 to regulate fork speed and pausing simultaneously.

**Figure 8:**
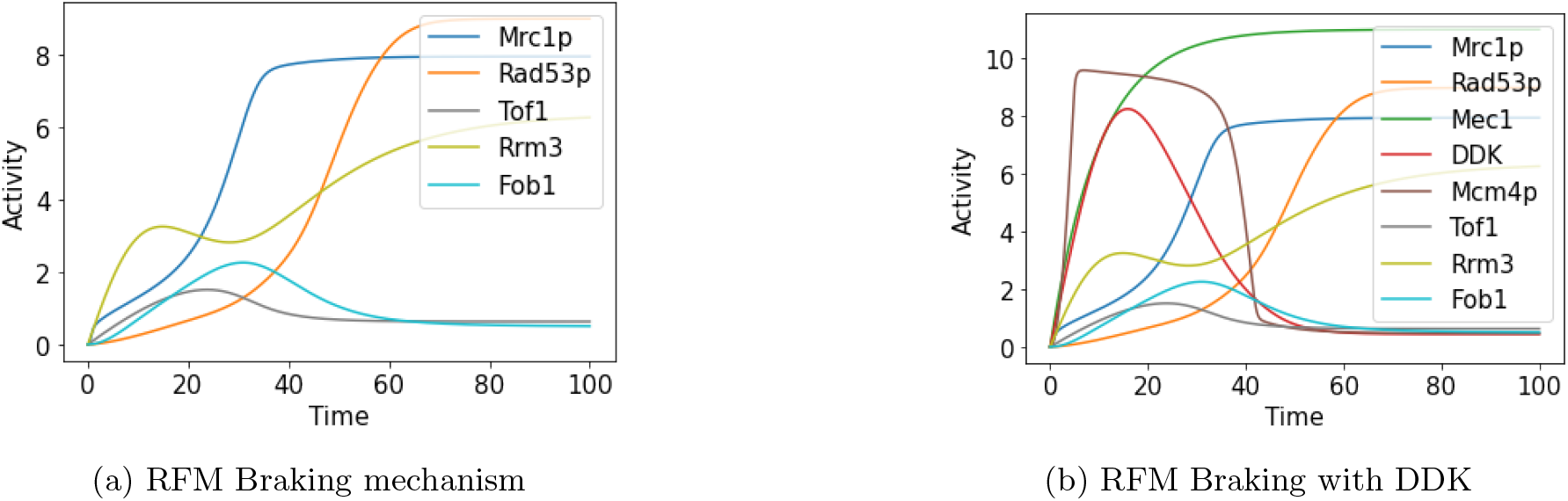
Simulation results of combined replication fork accelerator and braking mechanism. (a) High Mrc1P inhibits Tof1 activity resulting in high Rrm3 activity which negatively regulate fork progression and low Fob1 activity which cannot remove fork barriers resulting in slowing down fork progresison. (b) Initially during low replication stress Mec1 signal, Rad53P is low, DDK is high promoting firing of origins. But during high replication stress Mec1 signal, Rad53P becomes high inhibiting DDK activity in turn suppresses Mcm4P inactivates origin firing which results in pausing of the fork.

**Figure 9:**
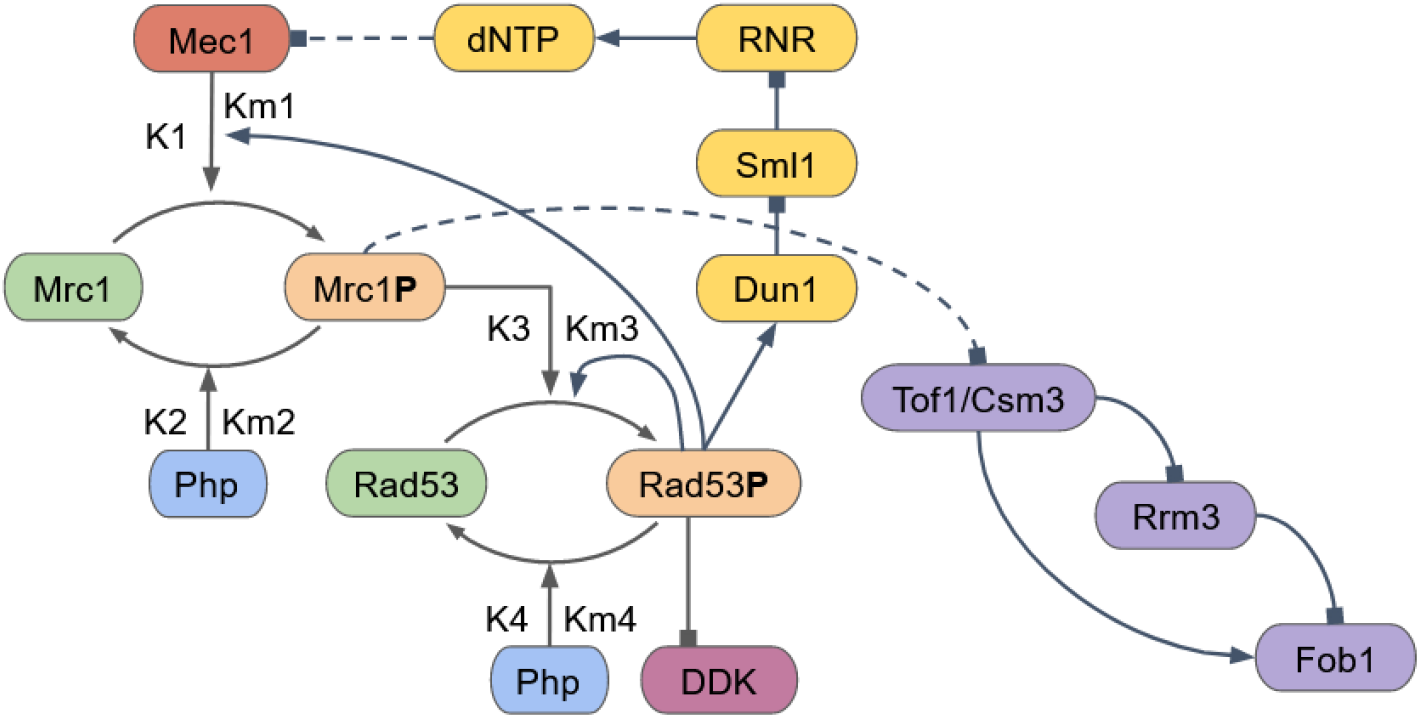
Replication Fork regulation using dNTP. It consists of both accelerator and braking mechanisms (shown in violet) along with dNTP regulation of replication stress signal (shown in yellow). Replication stress signal Mec1 is kept low when sufficient amount of dNTP is available during DNA replication. Low dNTP levels trigger stress signal Mec1 which results in slowing down of fork progression and inhibits formation of replication forks by suppressing firing of replication origins to prevent further DNA replication. When dNTP levels (shown in black) are high, Mec1 (shown in green) signal is low and vice versa.

**Figure 10:**
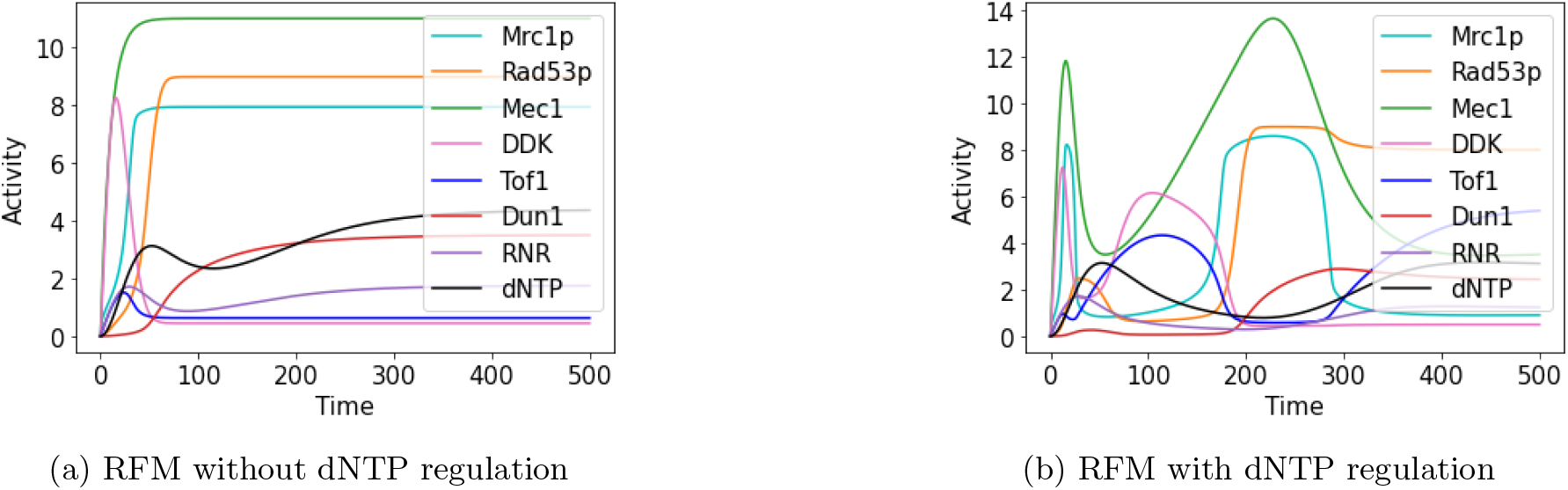
Simulation results of replication fork regulation using dNTP. (a) dNTP pathway does not regulate replication stress signal Mec1 and dynamics quickly reaches steady state. (b) dNTP negatively regulates replication stress signal sensor protein Mec1 which gives rise to a non-linear set of behavior of the complex system for some time before settling to a steady state. Complete list of all the parameters and rate constants is available in supplementary material.

**Figure 11:**
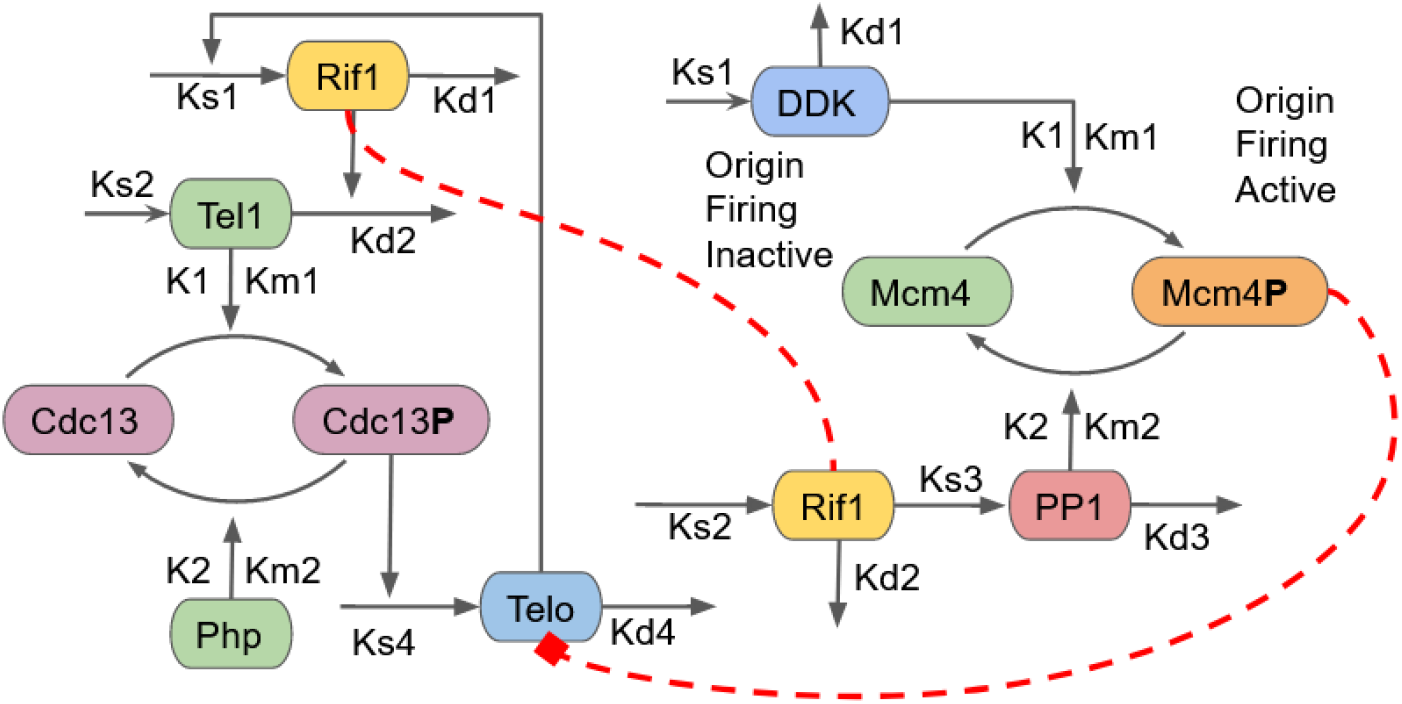
Telomere Length Homeostasis model with replication fork Mcm4 component along with DDK kinase. Interaction (red dash line) between Rif1 in telomeric DNA and nearby incoming replication fork is yet to be understood properly. Also, assumed Mcm4P interaction (red dash line) with telomerase (Telo) need to be validated experimentally.

**Figure 12:**
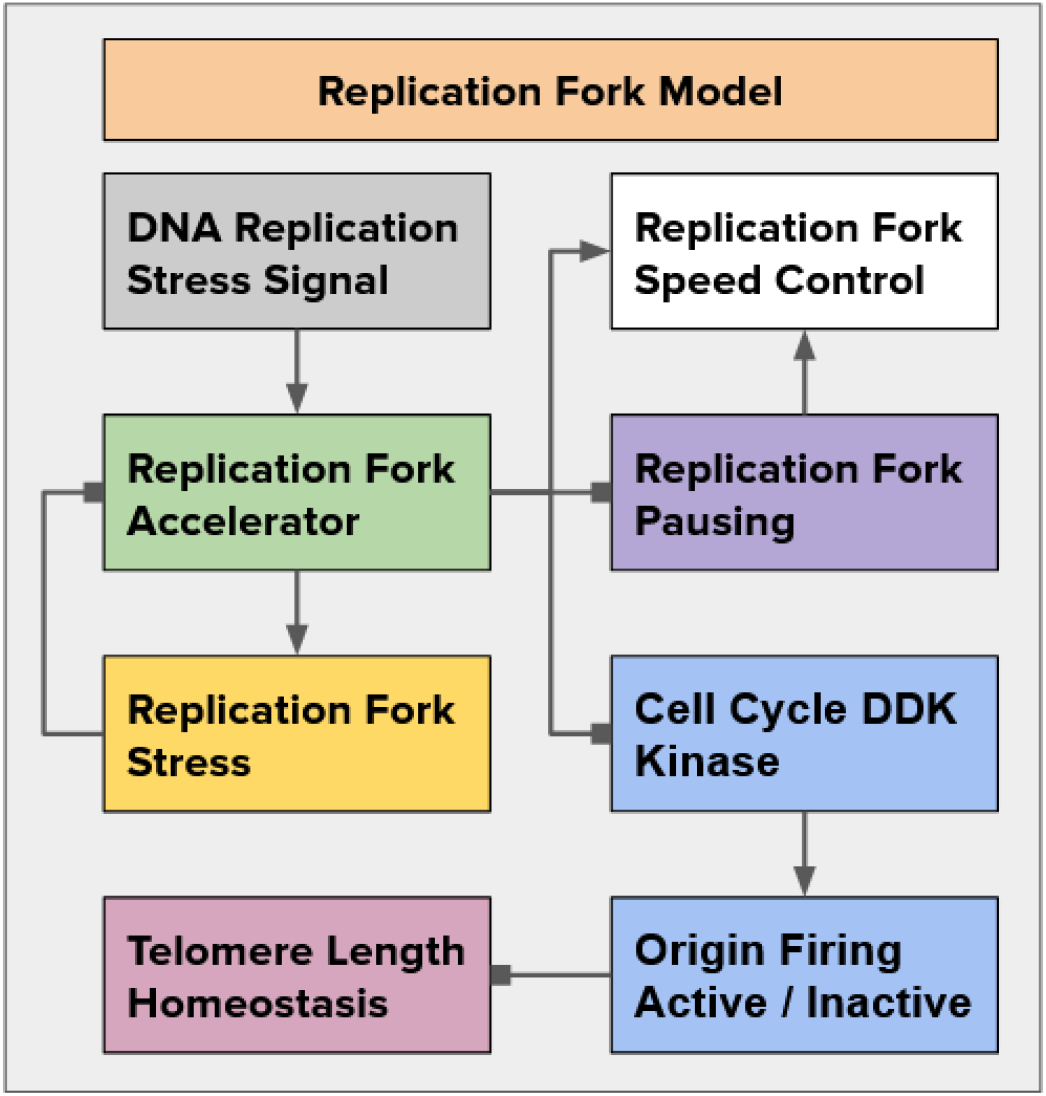
Proposed block diagram of complete replication fork model (CRFM).

### Replication Fork regulation using dNTP

In S phase replication checkpoints of S. cerevisiae, two signaling pathways converge at Mec1/Ddc2, which phosphorylates Rad53 with the help of Rad9 (DNA damage) or Mrc1 (replication stress). Phosphorylated Rad53 then activates Dun1, which, in turn, inactivates three repressors (Sml1, Crt1, and Dif1) of RNR. RNR generates dNTPs (deoxyribonucleotide triphosphate) from NDPs (Nucleoside triphosphate), which are then converted to dNTPs by nucleoside diphosphate kinases. Higher dNTP concentrations facilitate DNA repair and replication fork restart [34]. During the normal S phase, when the dNTP pool is low, Mec1 gets activated resultsing in pausing replication fork during DNA replication [11]. System of equations for model is given by:

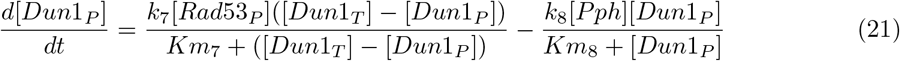

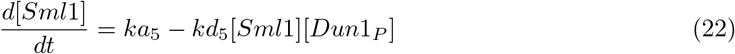

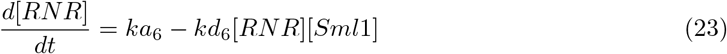

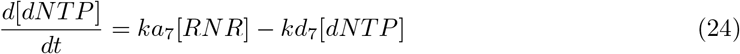

In equation 21, *k*_7_ is [*Rad*53_*P*_] mediated phosphorylation rate of [*Dun*1_*P*_], *k*_8_ is [*Pph*] phosphatase mediated dephosphorylation of [*Dun*1_*P*_], *Km*_7_ and *Km*_8_ are Michaelis constant and total concentration of [Dun1] is conserved [*Dun*1] + [*Dun*1_*P*_] = [*Dun*1_*T*_]. In equation 22, *ka*_5_ is [Sml1] production rate and *kd*_5_ is [*Dun*1_*P*_] mediated degradation rate of [Sml1]. In equation 23, *ka*_6_ is [RNR] production rate and *kd*_6_ is [Sml1] mediated [RNR] degradation rate. Further in equation 24, *ka*_7_ is [RNR] mediated [dNTP] production rate and *kd*_7_ is [dNTP] degradation rate.

### Replication Fork near Telomeres

The replication origins at internal Y’ elements can fire in the early mid-S phase, while ARSs at the terminal Y’ elements were confirmed to fire late. Telomeric pause strength was dependent upon telomere length and did not require the presence of a variety of factors implicated in telomere metabolism and known to cause telomere shortening [18]. In wild-type *S.cerevisiae,* replication forks slowed during their passage through telomeric C1–3A/TG1–3 tracts [14]. Rif1 Controls DNA Replication Timing in yeast through the PP1 Phosphatase [20]. The checkpoint regulatory proteins Tof1 and Mrc1 interact directly with the DNA replication machinery in *S.cerevisiae.* When hydroxyurea blocks chromosomal replication, this assembly forms a stable pausing structure and anchor subsequent DNA repair events. Also, these fork progression/protection complex proteins Mrc1 and Tof1 travel with the replication fork [15]. DDK plays a specific role in DNA double-strand breaks (DSBs) [23] and RRM3 promote replication fork progression by removing fork barrier proteins [4]. Arrival of replication fork near telomeric regions results in slow down of fork progression. Near telomeres more no. of Rif1 proteins is present though telomere length regulation by Rif1 is different from Rif1 regulating firing of replication origins in telomeric and non telomeric regions [27].

Replication fork indirectly regulates telomere elongation and shortening. Assuming telomerase travels with the replicaiton fork. DDK kinase is required to activate telomeric replication fork by firing subtelomeric origins. Based on the simulation results we found that high DDK kinase phosphorylates replication fork factor Mcm4p 2(b) which is needed to fire origins. High amount of Mcm4p inhibits activity of telomerase resulting in shortening of telomeres and reduces negative effect due to less telomere bound Rif1 protein as shown in figure 2(a). Later when the replication fork factor Mcm4p activity is reduced, it cannot inhibit telomerase and telomerase can come and elongates telomeres.

Mathematically Rif1, Telo, and Mcm4p are the key variables relating replication fork with telomeres regulation based on the assumption that Mcm4 inhibits telomerase. *Ks*_1_ and *Kd*_4_ are key parameters playing a role in telomere regulation by replication fork. Still the assumption need to be experimentally tested and validated for concrete understanding of the nature of interaction between telomeric Rif1 proteins and incoming replication fork.

## Conclusions

In this research work, we adopted a modular approach and initially developed different components of the replication fork model independently and later connected them to get the global picture. Initially, we modeled the S phase DNA replication checkpoint as a positive feedback loop. It gives rise to bi-stable behavior, as seen from the phase plane and bifurcation analysis indicating replication fork progression makes a transition from faster to slower dynamics during replication stress encountered during DNA replication. We call this behavior a replication fork accelerator. Next, we considered the role of crucial cell cycle factor CDK in licensing origins and DDK kinase in activating replication origins and showed how inhibition of DDK by Rad53 inhibited origins from firing. Later, we developed a simple replication fork pausing mechanistic circuits as a coherent feed-forward loop based on the recent finding of the ‘‘molecular brake” concept and calls this replication fork pausing behavior. Then we connected the replication fork accelerator and pausing together using Mrc1 dynamic interaction with MTC (Mrc1, Tof1, Csm3) complex and obtained replication fork accelerator pausing model. Our simulation results predicts that the replication fork’s acceleration and pausing are two sides of the same coin. Lastly, we considered the replication fork stress pathway and the role of dNTP depletion in regulating Mec1, which senses stress signals. Finally, we connected the replication fork model with the telomere length homeostasis model to understand the role of DNA replication and origin firing in telomere length regulation. We modeled the replication fork system using ODEs and analyzed using dynamical systems theory to get insights into the fork dynamics. For further understanding, we need to develop a mathematical equation to explain the importance of replication fork speed during DNA replication, which connects with origin firing rate, fork density and gives more insights on how replication fork regulates telomere length in near future.

## Future Work

Reduced replication fork speed promotes pancreatic endocrine differentiation and controls graft size. Reduced replication fork speed during differentiation improves the stability of insulin expression. Important for pancreatic endocrine differentiation [31]. Mouse embryonic stem cells (ESCs) utilizing a unique Filia-Flopped protein complex dependent mechanism to efficiently promote the restart of stalled replication forks and maintains genomic stability. Mouse ESCs are superior to differentiated cells in stabilizing and restarting the stalled replication forks [36]. So it will be exciting and important to identify the replication fork velocity roles with other metabolic processes.

## Acknowledgement

Research work design, modeling, and simulation carried independently by Ghanendra Singh along with the M.Tech. Thesis research work at the Center for Computational Biology, IIIT Delhi. Thanks to Dr. Sriram K. for providing valuable suggestions and feedback on the manuscript.

## Notes

https://github.com/Ghanendra19213/Telomere_End_Replication_Problem/tree/main/Replication%20Fork%20Model

### Competing Interest Statement

The authors have declared no competing interest.

