## Supplementary Material for "Mechanistic Model of Replication Fork Progression in Yeast"

### Mechanistic Model of Replication Fork Progression in Yeast Supplementary Material

Ghanendra Singh[1]

[1]Center for Computational Biology IIIT Delhi

#### Model Parameters

##### Replication Fork Accelerator Model Parameters

| Parameters | Description | Value |
|---|---|---|
| $k_1$ | [Mrc1] activation rate | 0.05 |
| $k_2$ | [Mrc1] deactivation rate | 0.03 |
| $k_3$ | [Rad53] activation rate | 0.03 |
| $k_4$ | [Rad53] deactivation rate | 0.02 |
| $Km_1$ | Michaelis constant | 1.0 |
| $Km_2$ | Michaelis constant | 1.0 |
| $Km_3$ | Michaelis constant | 1.0 |
| $Km_4$ | Michaelis constant | 1.0 |
| $k_a$ | [Mec1] production rate | 1.1 |
| $k_d$ | [Mec1] degradation rate | 0.1 |
| $k_{a2}$ | [DDK] production rate | 0.8 |
| $k_{d2}$ | [DDK] degradation rate | 0.2 |

Table 1: Replication Fork Accelerator Model

##### Cell Cycle Parameters

$a_1 = 0.02$, $a_2 = 3$, $a_3 = 3$, $b_1 = 3$, $b_2 = 1$, $K_1 = 0.5$, $K_2 = 0.5$, $K_3 = 0.5$, $n_1 = 8$, $n_2 = 8$, $n_3 = 8$

#### Replication Fork Pausing Parameters

| Parameters | Description | Value |
|---|---|---|
| $kp_1$ | [Tof1] production rate | 0.1 |
| $bd_1$ | [Tof1] degradation rate | 0.02 |
| $kp_2$ | [Rrm3] production rate | 0.4 |
| $bd_2$ | [Rrm3] degradation rate | 0.1 |
| $kp_3$ | [Fob1] production rate | 0.2 |
| $bd_3$ | [Fob1] degradation rate | 0.04 |

Table 2: Replication Fork Pausing Parameters

#### Replication Fork dNTP Regulation Parameters

| Parameters | Description | Value |
|---|---|---|
| $k_7$ | [Dun1] activation rate | 0.02 |
| $k_8$ | [Dun1] deactivation rate | 0.02 |
| $Km_7$ | Michaelis constant | 1.0 |
| $Km_8$ | Michaelis constant | 1.0 |
| $ka_5$ | [Sml1] production rate | 0.1 |
| $kd_5$ | [Sml1] degradation rate | 0.01 |
| $ka_6$ | [RNR] production rate | 0.1 |
| $kd_6$ | [RNR] degradation rate | 0.02 |
| $ka_7$ | [dNTP] production rate | 0.1 |
| $kd_7$ | [dNTP] degradation rate | 0.04 |

Table 3: Replication Fork dNTP Parameters

#### Initial values

[Mec1] =(1-10), [Mrc1T]=10, [Rad53T]=10, [Php]=10, [Dun1T]=10. All Initial values (t=0) = 0.
